# Loss of thymocyte competition underlies the tumor suppressive functions of the E2a transcription factor in T lymphocyte acute lymphoblastic leukemia

**DOI:** 10.1101/2023.04.23.537993

**Authors:** Geoffrey Parriott, Emma Hegermiller, Rosemary E. Morman, Cameron Frank, Caner Saygin, Wendy Stock, Elizabeth T. Bartom, Barbara L. Kee

## Abstract

T lymphocyte acute lymphoblastic leukemia (T-ALL) is frequently associated with increased expression of the E protein transcription factor inhibitors TAL1 and LYL1. In mouse models, ectopic expression of Tal1 or Lyl1 in T cell progenitors or inactivation of E2a, is sufficient to predispose mice to develop T-ALL. How E2a suppresses thymocyte transformation is currently unknown. Here, we show that early deletion of *E2a*, prior to the DN3 stage, was required for robust leukemogenesis and was associated with alterations in thymus cellularity, T cell differentiation, and gene expression in immature CD4+CD8+ thymocytes. Introduction of wild-type thymocytes into mice with early deletion of *E2a* prevented leukemogenesis, or delayed disease onset, and impacted the expression of multiple genes associated with transformation and genome instability. Our data indicate that E2a suppresses leukemogenesis by promoting T cell development and enforcing inter-thymocyte competition, a mechanism that is emerging as a safeguard against thymocyte transformation. These studies have implications for understanding how multiple essential regulators of T cell development suppress T-ALL and support the hypothesis that thymus cellularity is a determinant of leukemogenesis.

---

Acute lymphoblastic leukemia is the most common form of pediatric cancer, with T lymphocytes (T-ALL) accounting for 10-15% of cases (Karrman and Johansson, 2017; Aifantis et al., 2008). T-ALL is a heterogeneous malignancy that arises from the transformation of T cell progenitors in the thymus. It is divided into subgroups based on specific mutations and gene expression profiles (Belver and Ferrando, 2016; Noronha et al., 2019), for example, mutations affecting *LYL1* and *LMO2* are associated with an early thymic progenitor subtype of T-ALL (ETP-ALL), while ectopic expression of TAL1, HOXA, or GATA3 is associated with more mature double-positive (DP)/Cortical like T-ALL (Ferrando and Look, 2003; Coustan-Smith et al., 2009). These subset specific mutations are found alongside mutations that are represented across all multiple subtypes of T-ALL, including mutations affecting *NOTCH1*, *MYC*, *CDKN2A*, and *EZH2* (Coustan-Smith et al., 2009; Noronha et al., 2019; Aifantis et al., 2008; Van Vlierberghe and Ferrando, 2012; O’Neil et al., 2006). However, the mechanisms by which these genetic alterations are acquired and how they contribute to the transformation process is largely unresolved. Understanding these mechanisms may provide valuable insights into more effective and less toxic therapies, as well as preventing relapse.

Many T-ALL subsets exhibit aberrant expression of proteins that affect the function of the E protein helix-loop-helix (bHLH) transcription factors encoded by the *Tcf3* (E2a) and *Tcf12* (Heb) genes (Parriott and Kee, 2022). The E proteins are critical for T cell development, with E2a playing important roles before and after the seeding of progenitors in the thymus (Bain et al., 1997; Yan et al., 1997; Barndt et al., 2000). In multipotent hematopoietic progenitors, where Heb is not highly expressed, E2a forms homodimers or heterodimers with the hematopoietic stem cell (HSC)-associated bHLH transcription factors Lyl1 and Tal1 (Wilson et al., 2010; Bouderlique et al., 2019). During lymphoid specification Tal1, which can repress lymphoid gene expression, decreases and E2a:Lyl1 dimers support the generation of thymic seeding progenitors (de Pooter et al., 2019; Zohren et al., 2012). As thymocytes commit to the T cell lineage, *Lyl1* is extinguished, and *Heb* increases, resulting in E2a:Heb dimers that support multiple stages of T cell development (Yui and Rothenberg, 2014; Anderson, 2006). Aberrant expression of TAL1 or LYL1 is found in over 70% of pediatric T-ALL patients (60% and 10% of cases, respectively) (Ferrando and Look, 2003). TAL1 is thought to induce leukemia through multiple mechanisms, including formation of a TAL1:E2A:LMO2 complex that drives expression of oncogenes like c-*MYB* and *GATA3* (Tan et al., 2019; Sanda et al., 2012). However, in mice, T cell progenitors forced to express of a form of Tal1 that prevents DNA binding by Tal1:E2a/Heb develop T-ALL-like disease, which is exacerbated by heterozygous deletion of E2a (O’Neil et al., 2004). These finding suggest that TAL1 and LYL1 may function by inhibiting E2A:HEB transcriptional activity. Consistent with this hypothesis, mice lacking *E2a* or ectopically expressing inhibitors of E protein DNA binding in thymocytes develop T-ALL (Kim et al., 1999; Morrow et al., 1999). Moreover, some cases of ETP-ALL and DP/Cortical T-ALL overexpress ID proteins, which are inhibitors of E protein DNA binding (Coustan-Smith et al., 2009). Therefore, E protein inhibition is a common feature of human T-ALL.

Recent studies have implicated inter-thymocyte competition as a mechanism preventing leukemogenesis (Paiva et al., 2018). In experimental settings where wild-type (WT) thymocytes develop without incoming thymus seeding progenitors, aging WT thymocytes maintain T cell production, called “thymus autonomy”, acquire somatic mutations, and progress to T-ALL (Martins et al., 2014). Leukemogenesis can be suppressed in thymus autonomy by restoring competition with young thymocytes. However, competition is less effective at suppressing transformation after 6-8 weeks of thymus autonomy suggesting that critical transforming events may be established by that time (Martins et al., 2014). Leukemias arising from thymus autonomy acquire *Notch1* mutations and express *Tal1* and *Lmo2*, suggesting that thymus autonomy creates a selective pressure for E protein inhibition (Martins et al., 2014; Weng et al., 2004; Reschly et al., 2006). Recent studies in Lmo2 transgenic mice indicate that compromised T cell differentiation also leads to a failure of competition and outgrowth, in this case, of ETP-ALL (Abdulla et al., 2023; McCormack et al., 2013). Here, we used multiple models of E2a deficiency to investigate the mechanisms underly the development of T-ALL and found that early deletion of *E2a*, in HSCs, more robustly predisposed mice to develop T-ALL than *E2a* deletion in DN3 thymocytes. Early *E2a* deletion was associated with a pronounced deficiency of CD4+CD8+ (DP) thymocytes and upregulated expression of *Notch1* and *Myc* mRNA and pathways associated cancer and genome instability in DP thymocytes. We demonstrate that leukemogenesis could be suppressed in these mice by placing pre-leukemic cells in an environment with a sufficient number of WT thymocytes to promote competition. Thus, E2a functions as a suppressor of leukemogenesis, at least in part, by supporting proper T cell development and inter-thymocyte competition. Our findings demonstrate that intrinsic mutations in hematopoietic progenitors can lead to T-ALL by impacting extrinsic mechanisms that purge cells with mutations that make them less competitive but primed for leukemogenesis in the absence of competition. These studies have implications not only for T-ALL arising from genomic alterations affecting E2A and other regulators of T cell development but also for T-ALL arising in the context of lymphocyte progenitor deficiency.

## RESULTS

### *E2a* deletion before the DN3 thymocyte stage is required for efficient leukemogenesis

The T-ALLs arising from *E2a^-/-^* mice have a CD4^+^CD8^+^ (DP) phenotype with variable T cell receptor expression, but they also have features of DN3 thymocytes including expression of the Notch1 target gene *Il2ra* (CD25) (Adler et al., 2003), which is usually downregulated during β-selection (Bain et al., 1997; Carr et al., 2022). *E2a^-/-^* mice have multiple defects in the generation of T cell progenitors, including a failure to generate lymphoid primed multipotent progenitors and their expression of the thymus homing chemokine Ccr9 (Dias et al., 2008; Krishnamoorthy et al., 2015; Yang et al., 2011), increased apoptosis in DN2 thymocytes (Kee et al., 2002), a failure to induced *Notch1* (Ikawa et al., 2006), and diversion to the innate lymphoid fates (Xu et al., 2013; Qian et al., 2019) Given these early requirements for E2a, we questioned whether loss of *E2a* at the DN3 stage would be sufficient to predispose mice to T-ALL. To test this, we created *Lck^Cre^E2a^f/f^*(LcKO) mice, where *E2a* is deleted at the DN3 stage, and *Vav^Cre^E2a^f/f^*(VcKO) mice, where *E2a* is deleted at the hematopoietic stem cell stage on a leukemia susceptible FVB/NJ background and then evaluated leukemia latency and penetrance. As expected, VcKO mice rapidly developed leukemia with a mean latency of 17.4 weeks and 100% penetrance (Fig. 1A, B). In contrast, only 4/13 (30.8%) LcKO mice developed T-ALL within the 52-week evaluation with an increased mean latency of 37.3 weeks (Fig. 1A, B). In LcKO mice, the *E2a^f^* allele was deleted in >70% of DN3 thymocytes and these cells had reduced E2a protein (Sup. Fig. 1A and B). Taken together, these data indicate that *E2a* deletion in HSCs results in an increased penetrance and reduced latency of leukemia compared to deletion in DN3 thymocytes.

**Fig. 1.**
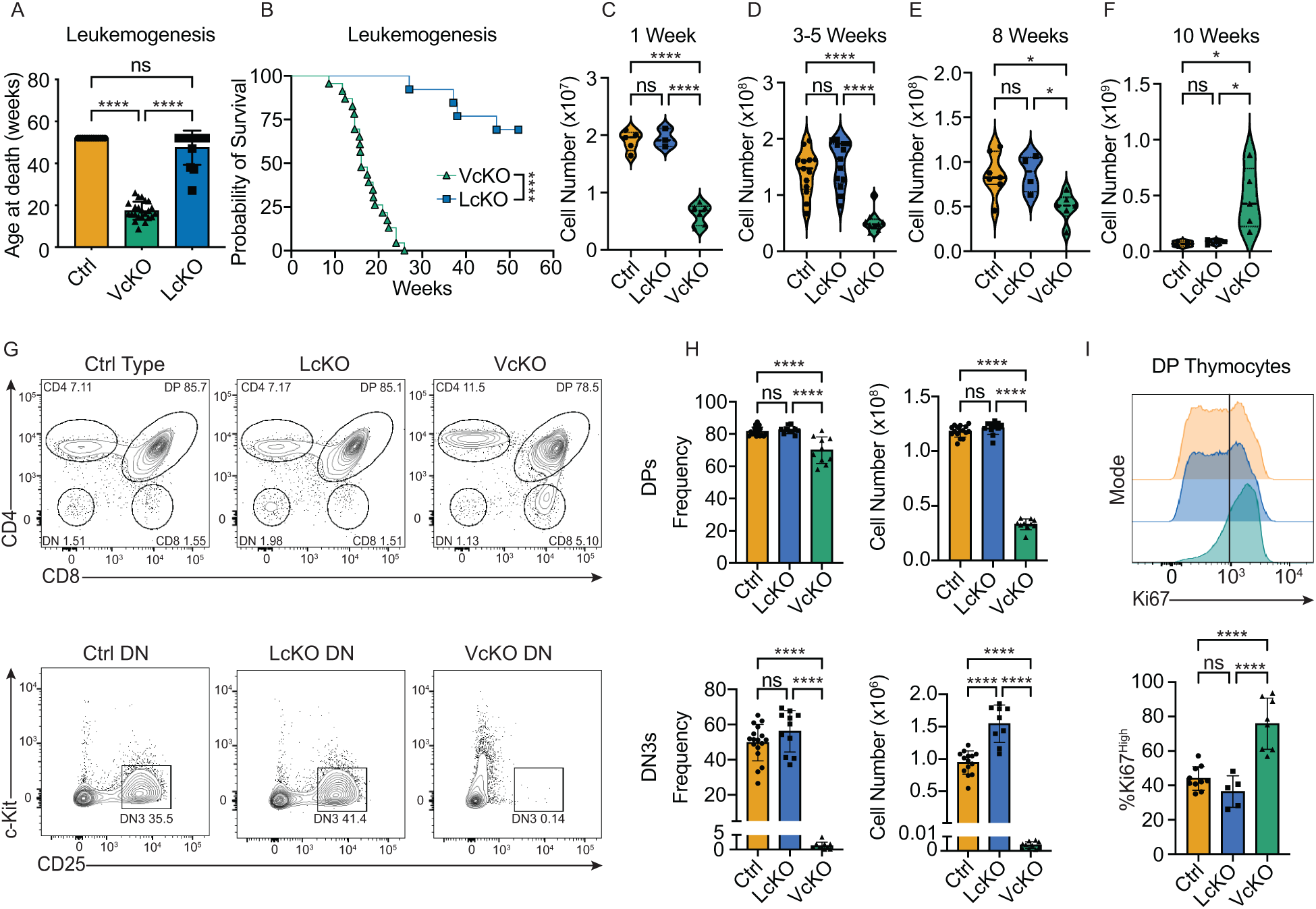
Early hematopoietic deletion of *E2a* predisposes mice to T-ALL and alters thymocyte development. (A) Age at death of VcKO, LcKO, and control mice throughout 52 weeks of study. (B) Kaplan-Meier curve of leukemia free survival for VcKO and LcKO mice. (C-F) Analysis of total number of cells in the thymus of VcKO, LcKO and Ctrl mice at (C) one week, (D) 3-5 weeks, (E) 8 weeks, and (F) 10 weeks of age. (G) Representative FACS plots showing CD4 and CD8 profiles gated on Lineage^-^ thymocytes (Top) or c-Kit and CD25 on Lineage^-^CD4^-^CD8^-^ thymocytes (Bottom). (H) Quantification of frequency and number of DP (Top) or DN3s (Bottom) thymocyte populations. (I) Representative FACS plots (top) and quantification of Ki67 expression (bottom) in DP thymocytes.

### Thymocyte deficiency requires early deletion of *E2a*

To gain insight into the differences between HSC and DN3 deletion of *E2a* we first examined thymocyte numbers. As expected, VcKO mice had a significant reduction of total thymocytes at 1 week, 3-5 weeks, and 8 weeks of age compared to Control (Ctrl) or LcKO mice (Fig. 1C-E). This difference was most significant in neonatal and young adult mice, but by 8 weeks the number of thymocytes from Ctrl and LcKO mice begin to contract, whereas VcKO thymocyte numbers stayed constant (Fig. 1C-E, Sup. Fig. 1C, D). By 10 weeks of age VcKO mice had more thymocytes than Ctrl or LcKO mice, and there was an expansion of VcKO thymocyte numbers compared to 8 weeks suggestive of the outgrowth of pre-leukemic or leukemic cells (Fig. 1F, Sup. Fig 1C). At 3-5 weeks of age, DN3 and DP cell frequencies and numbers were reduced in VcKO mice (Fig. 1G, H). This was not the case for LcKO mice, which had normal frequencies of DN3 and DP thymocytes (Fig. 1G, H). Analysis of Ki67, a protein expressed in all cycling cells, revealed that despite their reduced frequency and number there were more Ki67^high^ DP thymocytes in VcKO mice than in Ctrl or LcKO mice (Fig. 1I). Our data indicate that deletion of *E2a* in HSCs, but not in DN3 cells, results in reduced DN3 and DP cell numbers in mice prior to 10 weeks of age.

### Oncogene transcription in preleukemic VcKO DP thymocytes

To investigate how deletion of *E2a* in HSC or DN3 impacts the transcriptional program of DP thymocytes we performed RNA-seq on these cells from mice at 4 weeks of age. Analysis of the transcriptomes of Ctrl, VcKO, and LcKO DPs revealed that LcKO DPs more closely resemble WT DPs than VcKO DPs (Fig. 2A). Indeed, there were 419 differentially expressed genes (DEGs) in LcKO compared to Ctrl DPs, while VcKO DPs had 1659 DEGs (Fig. 2B). 744 DEGs upregulated in VcKO DPs showed no change in LcKO DPs, and 626 downregulated DEGs in VcKO DPs showed no change in LcKO DPs (Fig. 2B). There were DEGs that were specific to LcKO DPs, as 61 and 108 genes were upregulated or downregulated, respectively, in LcKO but not VcKO DP thymocytes (Fig. 2B). KEGG pathway analysis on the DEGs unique to VcKO DPs revealed an enrichment of genes in the Notch pathway, Jak/Stat signaling, chemokine signaling, cancer pathways, and several metabolism pathways (Fig. 2C). Gene set enrichment analysis (GSEA) confirmed enrichment of the Wnt_β_catenin, Myc, Notch and TNFA_via_NFκB pathway genes in VcKO DPs (Fig. 2D). Notably, only VcKO DPs had increased *Notch1* mRNA, and they had a substantially greater increase in the *Notch1* target *Il2ra* than LcKO DPs (Fig. 2E). Consistent with Myc pathway enrichment, only VcKO DPs, but not LcKO DPs, had a significant increase in *Myc* mRNA. Notch1 induces *Myc* expression in T-ALLs (Weng et al., 2006), and therefore increased *Notch1* expression could be associated with enrichment of the *Myc* pathway. Wnt pathway member *Lef1* was also increased in VcKO DPs (Fig. 2E). Despite the increased expression of *Notch1* mRNA, mRNA variants affecting exon 34, encoding the PEST domain of NOTCH1, which are prevalent and monoclonal in *E2a^-/-^* leukemias (Reschly et al., 2006), were rare in 4-week-old VcKO DPs and unlikely to account for the increased expression of this gene (Sup. Fig. 2A).

**Fig. 2.**
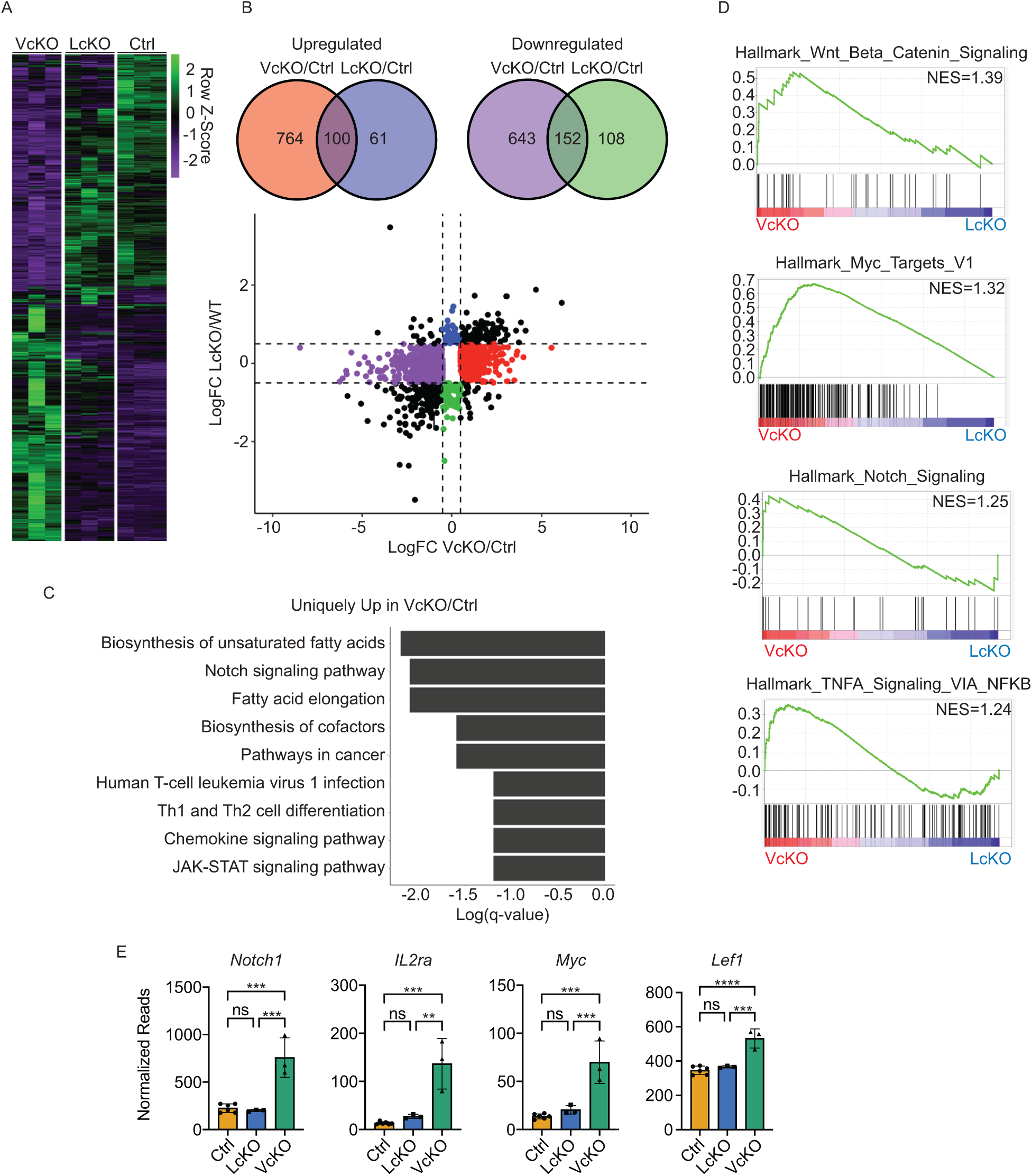
HSC-, but not DN3-deletion of *E2a* alters oncogenic gene expression in DP thymocytes from 4-week-old mice. (A) Heatmap of genes DEGs in either VcKO or LcKO DPs compared to Ctrl DPs. (B) Venn diagram showing number of DEGs between VcKO vs Ctrl DPs and LcKO vs Ctrl DPs (Top) and graph of LogFC over control for all DEGS in VcKO and LcKO DPs (Bottom). (C) Graph of log(q-value) of relevant KEGG pathways analysis of genes uniquely upregulated in VcKO vs Ctrl DPs. (D) GSEA plots showing pathways enriched in VcKO DPs compared to LcKO DPs. (E) Normalized RNA-sequencing reads showing altered gene expression between VcKO, LcKO and Ctrl DPs.

### Increased oncogene transcription in 8-week-old VcKO thymocytes

By 8 weeks of age, there were noticeable phenotypic differences in VcKO thymocytes compared to 3–5-week-old VcKO thymocytes including a reduced frequency of DP thymocytes with heterogeneous expression of CD4 or CD8 and an increased frequency of CD25^+^ cells (Fig. 3A-B), suggesting increased Notch1 activation or increased cytokine signaling (Adler et al., 2003). Surprisingly, the frequency of Ki67^High^ cells was not different in VcKO DPs from at 3-5 week or 8-week-old mice (Fig. 3C). To gain a better understanding of how the transcriptome of DPs changes as VcKO mice age, we performed RNA-seq on DPs from 8-week-old VcKO and Ctrl mice. There were 4,033 DEGs between 8-week VcKO and Ctrl DPs, a substantial increase compared to 1,637 DEGs in 4-week-old mice (Fig. 3D, Fig 2B). Of the 8-week DEGs, 1392 were dysregulated in both 4- and 8-week-old VcKO DPs (Fig. 3D, E). Many of the DEGs that increased in 4-week VcKO DPs were further upregulated in 8-week VcKO DPs, and the same trend was observed in DEGs that decreased in 4-week VcKO DPs, suggesting that exacerbated dysregulation of these gene may be associated with the emergence of pre-leukemic or leukemic cells (Fig. 2D, E). Many of the KEGG pathways enriched in 4-week DPs were also enriched in 8-week VcKO DPs, including Notch1 signaling, cancer associated pathways, and metabolic pathways (Sup. Fig. 2B). *Notch1* mRNA was expressed similarly in 4- and 8-week VcKO DPs but *Fbxw7* mRNA, encoding a ubiquitin ligase that targets Notch1 for degradation (Crusio et al., 2010), was decreased, which could promote the stability of activated Notch1 (Fig. 3F). However, one of the 8-week-old VcKO mouse showed multiple reads with *Notch1* exons 34 mutation indicating that oncogenic mutations in *Notch1* can occur by this stage of development (Sup. Fig. 2A). KEGG pathway analysis of genes uniquely dysregulated in 8-week VcKO DPs showed an increase in DNA replication and cell cycle associated pathways (Fig. 3G), consistent with the expansion of thymocytes seen in VcKO mice that begins at this time point (Fig. 1B). Notably, pathways indicative of genomic instability were also increased including Mismatch repair, Nucleotide excision repair, and the p53 pathway (Fig. 3G). Indeed, *Trp53* mRNA, encoding p53, is increased in 4-week and more so in 8-week VcKO mice (Sup. Fig. 2C). The p53 repressed gene *Tet2*, a guardian of genome stability, was nearly fully abrogated in 8-week VcKO DPs (Fig. 3F) (Laptenko and Prives, 2017). Despite the increase in genomic instability pathways, the apoptosis, senescence, and autophagy pathways were decreased in 8-week DPs, indicating that stress-associated pathways that lead to cell death are repressed (Fig. 3H).

**Fig. 3.**
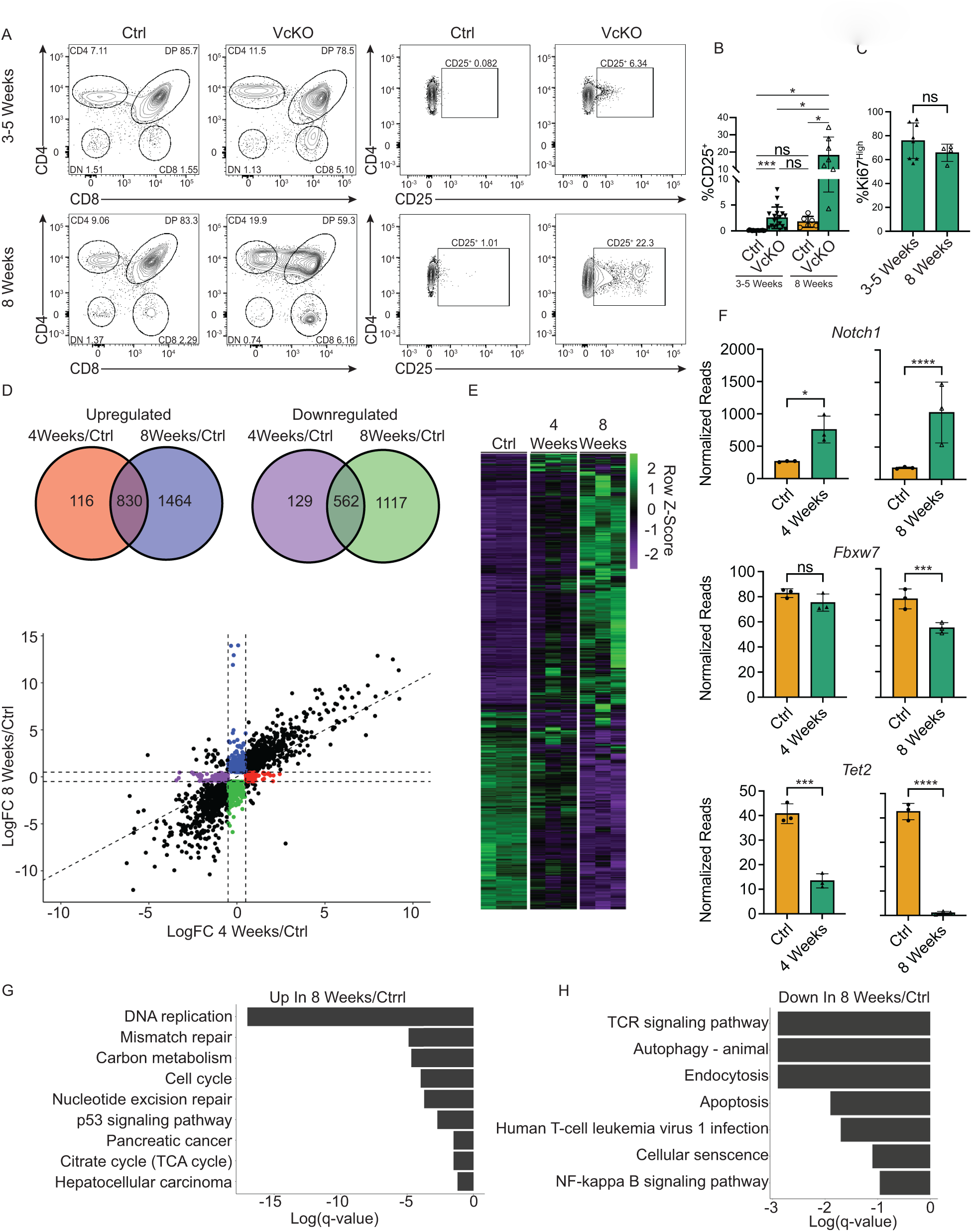
VcKO DP thymocytes show increased gene dysregulation in 8-week-old DPs compared to 3–5-week-old DPs. (A) Representative FACS plots showing CD4 and CD8 on lineage^-^ thymocytes (Left) and CD25 on DP thymocytes (Right) from 3–5-week (Top) or 8-week (Bottom) VcKO and Ctrl mice. (B) Frequency of CD25^+^ DPs from VcKO and Ctrl mice at the indicated age. Significance determined using Brown-Forscythe and Welch ANOVA. (D) Percent Ki67^High^ DPs in VcKO and Ctrl mice at the indicated age. (D) Venn diagram showing number of DEGs between VcKO DPs vs Ctrl DPs from 4 week or 8-week-old mice (Top) and graph of LogFC over Ctrl for all DEGS in 4 week and 8 week DPs (Bottom). (E) Heatmap of all DEGs in DPs vs Ctrl in 4 week or 8-week-old mice. (F) Normalized RNA-sequencing read counts showing altered gene expression between DPs vs Ctrl at 4 weeks and 8 weeks. Significance determined using EdgeR (see methods). (G-H) Graph of log(q-value) of relevant KEGG pathways analysis of genes uniquely (G) upregulated or (H) downregulated in 8 Week VcKO vs Ctrl DPs.

### Inter-thymocyte competition suppresses T-ALL development from VcKO cells

Our data indicate that deletion of *E2a* in HSCs results in profound alterations in T cell development and promotes a transcriptome marked by increased expression of *Notch1* and *Myc*, among other transformation-associated genes. Recent studies suggest that cell extrinsic mechanisms can contribute to thymocyte transformation and may cooperate with inhibition of E2a to drive leukemogenesis (Martins et al., 2014; Abdulla et al., 2023). In experimental models where WT thymi were deprived of incoming progenitors, thymocytes acquire a stem-like phenotype and eventually transform. Similar non-cell autonomous pathways have been shown in mice with defective generation of DP thymocytes (Abdulla et al., 2023). Given the multiple requirements for E2a in the generation of thymic seeding progenitors, and it’s role in promoting the DN2 to DN3 transition (Xu et al., 2013), we considered the possibility that VcKO thymocytes undergo transformation, at least in part, due to a failure of thymocyte competition. To investigate this, we took an approach that allowed a small number of WT hematopoietic progenitors to reconstitute VcKO mice. Three- to 6-week-old VcKO mice were sublethally irradiated and transplanted with T cell depleted WT (FVB/NJ CD45.2^+^) congenic bone marrow (BM) progenitors. Analysis of the VcKO thymi 4 weeks post-transplant showed that the CD4 by CD8 profile was similar to WT thymi, indicating that WT cells were able to successfully engraft and develop in the presence of VcKO cells (Fig. 4A). The total number of DPs was partially rescued in chimeric mice, although total thymocyte numbers were not different from VcKO mice (Fig. 4B). CD4 and CD8 single positive cells were also restored to near WT frequencies and numbers (Fig. 4B). CD45.1^+^ VcKO thymocytes could still be detected in the chimeras, although at reduced numbers as expected if these cells are not fit for competition with the transplanted WT cells. Thus, BM transplantation allowed for the introduction of WT T cell progenitors that can provide competition in VcKO mice.

**Fig. 4.**
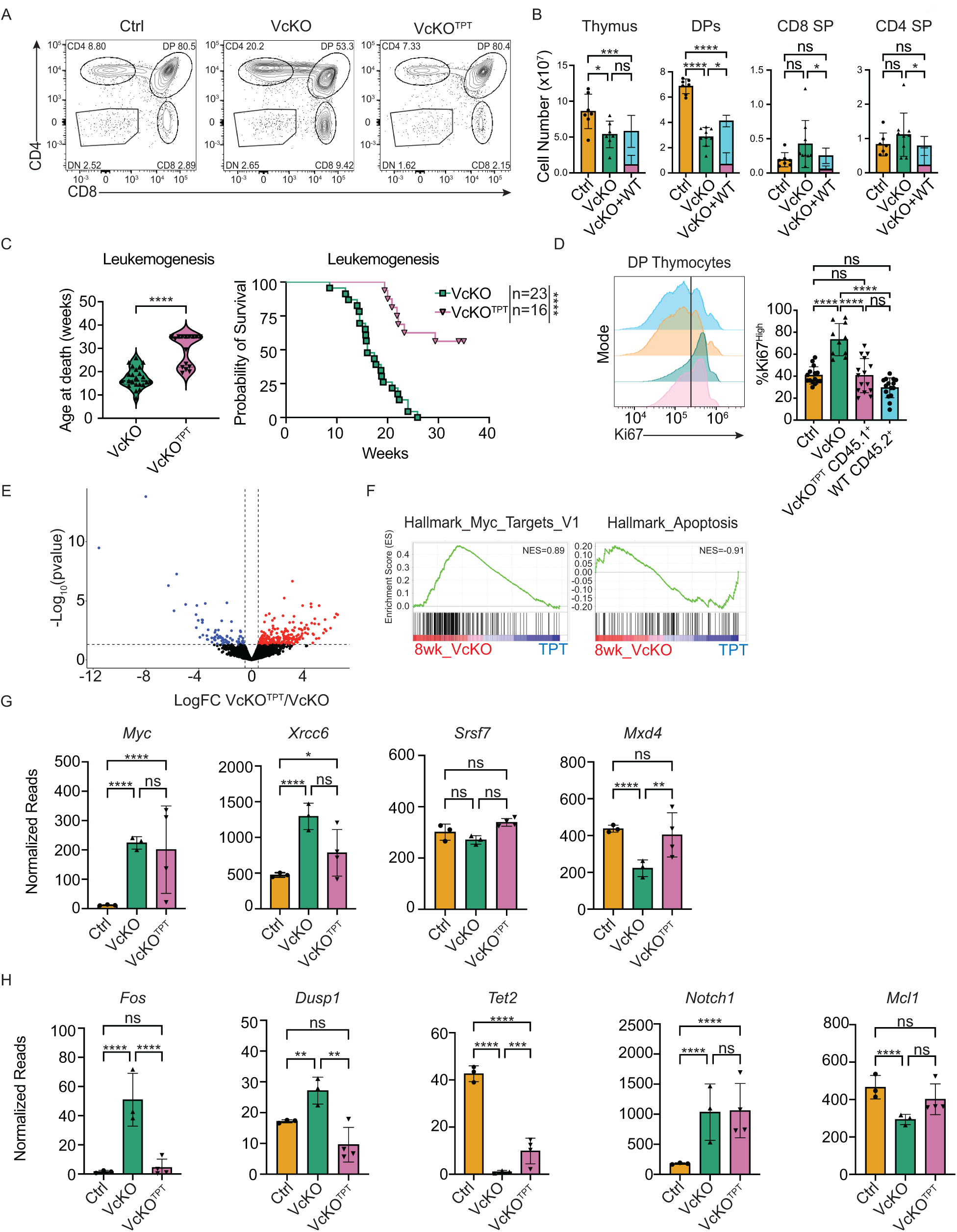
Wild-type thymocytes compete with VcKO thymocytes to limit leukemogenesis and alter DP gene expression. (A) Representative FACS plot showing CD4 and CD8 in the thymus of VcKO, VcKO+WT chimeras, and Ctrl mice. (B) Number of the indicated cell type from VcKO, VcKO+WT chimeras, and Ctrl mice. Cyan color indicates the total number of indicated populations. Pink color indicates the relative contribution of CD45.1^+^ (host-derived) cells. Statistics for this histogram are for total cell number. (C) Age at death of (Left) and Kaplan-Meier curve (Right) for VcKO and VcKO+WT chimeric mice. (D) Representative FACS histograms (Left) and quantification (Right) of Ki67 in VcKO, CD45.1^+^ (host derived) VcKO+WT, CD45.2^+^ (donor derived) in VcKO+WT chimeras, or Ctrl DPs. (E) Volcano plot showing significantly DEGs (p<0.05 and LogFC ≥ 0.5) between CD45.1+ VcKO^TPT^ and VcKO DPs. (F) GSEA showing pathways enriched in VcKO or VcKO^TPT^ DPs. (G) Normalized RNA-sequencing read counts showing expression of Myc pathways genes and (H) selected genes of interest in VcKO, VcKO^TPT^, and Ctrl DPs. Significance determined using EdgeR (see methods).

We monitored a cohort of VcKO+WT chimeric mice for 35 weeks to assess leukemia incidence and latency. VcKO+WT BM mice lived significantly longer than non-transplanted VcKO mice, with 9/16 mice surviving until 35 weeks (Fig. 4C). Analysis of a cohort of chimeric mice 4 weeks post-transplant revealed that Ki67^High^ DPs were significantly reduced in frequency among CD45.1+ VcKO (VcKO^TPT^) DPs in transplanted mice (Fig. 4D). To understand how competition affected the transcriptome of VcKO^TPT^ DPs, we performed bulk RNA-sequencing on CD45.1^+^ DP thymocytes from VcKO+WT chimeric mice 4 weeks after transplantation in comparison with age matched VcKO and WT DPs. We found substantial heterogeneity in the transcriptome of VcKO^TPT^ DPs but there were 351 DEGs between VcKO^TPT^ and VcKO DP thymocytes (Fig. 4E and S3A). Despite the heterogeneity, GSEA showed an enrichment of Myc pathways in VcKO DPs compared to VcKO^TPT^ (Fig. 4F). *Myc* mRNA was heterogeneous with two VcKO^TPT^ samples showing higher, and two showing lower, expression (Fig. 4G). *Xrcc6* mRNA was among the Myc targets that showed lower expression in 3 of 4 VcKO^TPT^ as compared to VcKO DPs (Fig. 4G). *Srsf7*, encoding a serine and arginine rich splicing factor, was a Myc target that was repressed slightly in VcKO DPs, but restored to WT levels in all VcKO^TPT^ samples, although this did not reach significance (Fig. 4G). Therefore, competition impacted expression of genes in the Myc pathway. There was also an enrichment for the hallmark apoptosis pathway in VcKO^TPT^ DPs with *Mxd4* mRNA, encoding for a Myc antagonist that inhibits leukemogenesis, topping the list of DEGs (Fig. 4G) (Hurlin et al., 1996) (Hu et al., 2022). Among the other notable DEGs were *Fos*, *Dusp1* and *Tet2*, which may explain one mechanism by which competition may regulate gene expression to promote survival (Fig. 4G). Interestingly, *Notch1* was not a DEG in VcKO^TPT^ as compared to VcKO DPs, but *Fbxw7* was increased (Fig. 4H). *Mcl1* was also increased in VcKO^TPT^ cells compared to VcKO, which was significantly different from Ctrl DP cells, which could be consistent with apoptosis being associated with transformation (Zhou et al., 1997).

### Competition promotes survival via exclusion of pre-leukemic cells

Analysis of leukemogenesis in VcKO+WT chimeras showed a bimodal response to transplantation. Of the 16 mice that we followed for 35 weeks 7 developed T-ALL within 30 weeks and 9 survived until the end of the study with no signs of leukemia (Fig. 5A, Fig. 4C). As the mice aged, we monitored chimerism in blood CD4^+^ T cells. VcKO+WT chimeras that survived >30 weeks (long-lived) showed robust chimerism in peripheral blood (>40% donor) at 4 weeks post-transplant. Mice that succumbed to T-ALL (short-lived) showed poor chimerism in the blood (<40% donor). With age, long-lived mice lost host cell contribution to peripheral blood CD4s progressively, while short lived mice continued to have a dominance of host cells (Fig. 5A). We then examined chimerism in the thymus in mice with <40% donor (Low chimerism) or >40% donor T cells in the periphery (High chimerism). In the thymus of VcKO+WT chimeric mice chimerism mirrored that of peripheral CD4^+^ T cells, with high chimerism mice showing >40% donor DP thymocytes, and low chimerism mice showing <40% donor DPs (Fig. 5B and C). This confirmed our use of peripheral CD4s as a read out of chimerism in the thymus. At 30 weeks post-transplant, long lived mice showed no host derived DP thymocytes in the remaining thymus tissue. Conversely, mice that succumbed to T-ALL at approximately 20 weeks post-transplant had no donor thymocytes. Therefore, in mice that survived until the end of the study, WT cells were able to efficiently colonize the thymus and eventually out compete pre-leukemic host cells, preventing leukemogenesis. In mice that became leukemic, donor cells colonize the thymus, but fail to out compete the host cells either because their numbers are insufficient or because transforming events have already occurred that increased the fitness of some host cells. To test if sufficient chimerism in the thymus is required to out compete pre-leukemic VcKO cells, we irradiated VcKO mice with a significantly lower dose of radiation (550 Rads) and transplanted congenically labeled WT BM cells. In these chimeric mice nearly all the thymocytes, including DPs, were of host origin (Sup. Fig. 3B and C). Further, all 550Rad VcKO+ BM mice fell into the poor chimerism group based on peripheral CD4 chimerism, and all 5 of them succumbed to T-ALL within 20 weeks (Fig. S3B, C). Together, these data indicate that there is a threshold of “fit” WT cells needed to provide sufficient competition to prevent leukemogenesis in VcKO mice.

**Fig. 5.**
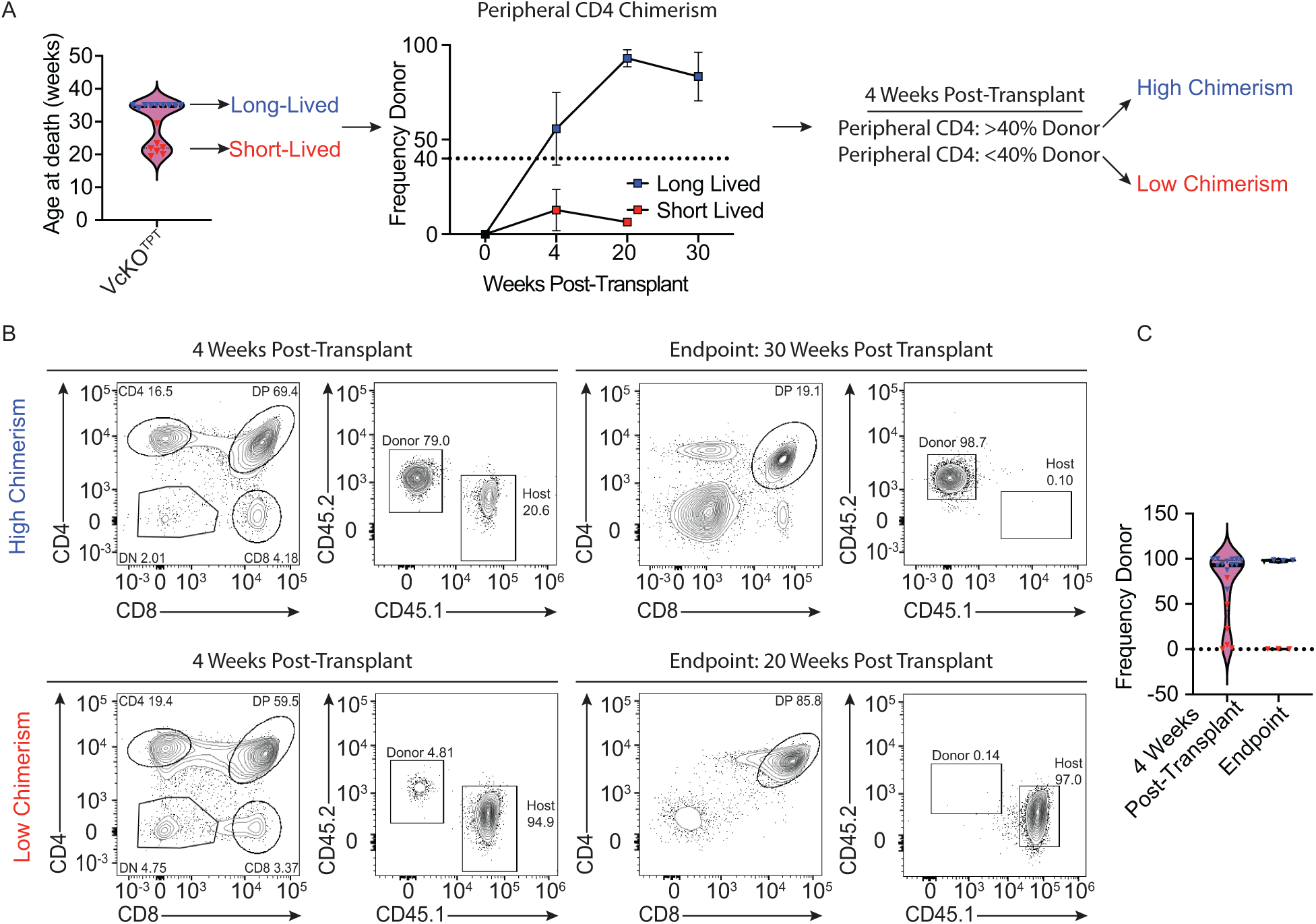
Leukemia latency in VcKO+WT chimeric mice is associated with WT competitor chimerism. (A) Violin plot (Left) showing age at death based on chimerism of VcKO+WT chimeric mice at 4 weeks post-transplant and their separation into high or low chimerism groups based on peripheral CD4 chimerism. The average frequency of donor cells at 4 weeks, 20 weeks or 30 weeks post-transplant is shown for high and low chimerism mice. Blue represents high chimerism and long-lived mice, red represents low chimerism and short-lived mice. (B) Representative FACS plots showing CD4 and CD8 profiles and CD45.1/CD45.2 chimerism of high chimerism mice (Top) and low chimerism mice (Bottom) at 4 weeks (Left) and at 22- or 30-weeks post-transplant (endpoint) (Right). (C) Frequency of donor thymocytes in high chimerism and low chimerism VcKO+WT chimeric mice at 4 weeks post-transplant and of long and short lived VcKO+WT chimeric mice at 20 or 30 weeks (endpoint).

### Discussion

Competition between fit and unfit cells in the thymus is emerging as an important mechanism to prevent leukemogenesis. Here we demonstrate that early hematopoietic deletion of *E2a* predisposes mice to T-ALL at least in part by its impact on T cell progenitor differentiation. Deletion of *E2a* in DN3 thymocytes did not impact DP thymocyte numbers and had a smaller impact on their gene expression profile than HSC deletion of *E2a* and rarely led to leukemic transformation. VcKO, but not LcKO DP, thymocytes show increased expression of *Notch1* and *Myc* mRNA and these pathways were further dysregulated by 8 weeks of age where some DP thymocytes show evidence of *Notch1* exon 34 mutations. Introduction of WT competitor cells delayed or prevented leukemogenesis depending on their ability to sufficiently engraft in the thymus and, presumably, the state of transformation of VcKO thymocytes at the time of transplantation. Competition impacts multiple genes in VcKO DP thymocytes that indicate repression of Myc function and dampening of genomic instability. Our data indicate that intrinsic mutations that impact T cell development can predispose to leukemic transformation by creating a thymic environment that fails to enforce the competition that removes unfit progenitors.

The need for thymocyte competition has been demonstrated in situations of complete thymus autonomy, whereas in our study we demonstrate that this mechanism is also important in a situation with reduced thymic input combined with impaired differentiation. These observations could be relevant in clonal hematopoiesis in which hematopoietic differentiation is skewed toward myelopoiesis rather than lymphopoiesis and could reduce the number of thymus seeding cells (Elias et al., 2017). Interestingly, repression of *Tet2* has been implicated in clonal hematopoiesis and is associated with T-ALL (Bensberg et al., 2021; Busque et al., 2012). We found decreased expression of *Tet2* in VcKO DP thymocytes that was exacerbated with age and its expression was partially restored in the presence of thymic competition. These observations suggest that *Tet2* could be a competition regulated gene that safeguards the genome of development thymocytes. Importantly, our data also suggest that regulators of MYC are impacted by competition.

CD2-Lmo2 transgenic mice develop ETP-like T-ALL and were recently shown to have deficient T cell differentiation from DN3 thymocytes, which was associated with apoptosis (Abdulla et al., 2023). Restoring thymocyte competition in these mice prevented transformation as did transgenic expression of Bcl2, which restored thymocyte differentiation. These observations suggest that induction of apoptosis could be a predisposing factor for T-ALL. *E2a^-/-^* mice have also have an early block in T cell development and undergo apoptosis (Kee et al., 2002). However, transgenic expression of Bcl2 did not restore differentiation in these mice indicating that apoptosis is not the driver of failed differentiation (Kee et al., 2002). Indeed, the arrested differentiation at the DN2 stage is mediated by an inability to properly express *Notch1* and repress *Gata3* (Xu et al., 2013; Ikawa et al., 2006), which may lead to diversion from the T cell fate (Xu et al., 2013; Qian et al., 2019). We note a prior study found that LcKO mice do not develop T-ALL (Pan et al., 2002). In contrast, our data demonstrate that on an FVB/NJ background *E2a* deletion in DN3s is sufficient to predispose mice to leukemia with incomplete penetrance and long latency suggesting that the non-competitive environment created by early deletion synergizes with E2a-deficiency to accelerate T-ALL.

It is notable that we did not find expansion of DP thymocytes with mutations in the PEST domain of Notch1 in 4-week-old VcKO mice, despite increased expression of *Notch1* mRNA. Nonetheless, there were a few reads identified with mutations that would affect the PEST domain indicating that, at this time point, there may be a polyclonal mutation spectrum in a subset of cells. By 8-weeks of age, we did observe multiple reads with the same mutation in two of 3 mice, although these mutations were still in only a small number of reads covering this region of exon 34. Notably, *Notch1* mutations are weak drivers of leukemogenesis and they do not increase Notch1 activity to the level seen in experimental models of activated Notch1 expression (Chiang et al., 2008). Therefore, in our leukemia model, Notch1 mutations might not be the drivers of the early increase in *Notch1* mRNA. The increase in *Notch1* mRNA may be driven by altered activity at the upstream enhancer that drives *Notch1* transcription in T cell progenitors (Kashiwagi et al., 2022), or by reduced expression of FBW7, which could prolong Notch1 activity (O’Neil et al., 2007; Thompson et al., 2007). Decreased FBW7 could also contribute to increased Myc either directly or indirectly through its effects on Notch1 degradation (Yada et al., 2004; Palomero et al., 2006; Weng et al., 2006). The Notch1-Myc axis is of particular interest in the VcKO+WT chimera setting where *Notch1* mRNA was not affected by introducing competition yet Myc pathway genes, and in some mice *Myc* itself, was decreased. There is a strong selection for Notch1-mediated regulation of Myc in T-ALL (Chiang et al., 2016), and our data suggest that this pathway may be kept in check by inter-thymocyte competition. It is not clear why some VcKO+WT chimera mice eventually succumb to T-ALL, however, Notch1 mutations may have already occurred in these mice that maintain high *Myc* expression despite the presence of competing thymocytes.

Taken together, our data indicate that the rapid onset of leukemogenesis in mice with an early deletion of *E2a* is mediated, in part, by E2a’s essential functions in promoting T cell development. This may be related to why multiple mouse models with altered T cell development also develop T-ALL. However, our data also suggest that reduced E protein function can synergize with a loss of thymocyte competition to accelerate the latency of leukemogenesis since *E2a*-deletion alone (in LcKO mice) is weakly oncogenic. These findings have significance for a broad range of T-ALL with altered E protein function, including those with increased LYL1, TAL1 or ID proteins, and suggest that reduced T cell differentiation may be a predisposing factor for leukemogenesis.

## METHODS AND MATERIALS

### Mice

VavCre mice (Jax #035670), LckCre (Jax, #003802), *E2a^F/F^* (Pan et al., 2002), and C57BL/6J mice were backcrossed onto an FVB/NJ background for 12 generations. The C57BL/6 backcrosses were selected for expression of CD45.2. Experimental mice were age-matched littermates and sex-matched whenever possible. Control (Ctrl) mice were *Cre^-^E2A^f/f^* or *Cre^-^E2A^f/+^*. All experiments were performed in accordance with the guidelines and approval of The University of Chicago Institutional Animal Care and Use Committee.

### Flow Cytometry

Tissues were harvested, and single cell suspensions were prepared. Samples were strained through 70μm filter and washed with ice-cold FACS + 1mM EDTA. Cells were treated with FC block (2.4G2, 1:200) and surface markers were stained for 25 minutes in the dark on ice. For some experiments, cells were stained with a lineage cocktail containing the following biotinylated antibodies at a 1:400 dilution: Ter119, CD11b, CD11c, DX5, GR1, B220, and CD19. For Ki67 analysis, cells were fixed and permeabilized using the FoxP3/Transcription factor staining buffer set (eBioscience Cat. #00-5523) according to manufacturer’s protocols. Fixed and permeabilized cells were stained with anti-Ki67-FITC (SolA15) at 1:500 dilution for 30 minutes at 4°C. For E2a analysis, cells were processed as noted in Ki67 analysis and stained with anti-E47 (G127-32) at 1:100 dilution for 30 minutes in the dark at 4°C. Cells were then stained with αMouse IgG Fab2-AlexaFluor 488 (Cell Signaling Technology Cat. #4408S) at 1:300 for 30 minutes in the dark at 4°C. Data were acquired on a LSRFortessa 4-15 (BD Biosciences), LSRFortessa X-20 (BD Biosciences), or NovoCyte Penteon (Agilent) flow cytometer in the Cytometry and Antibody Technology Facility at the University of Chicago. Data were analyzed using FlowJo v10.8.1 software.

### Transplantation assays

Bone marrow was isolated from the femurs and tibias of wild type FVB/NJ mice and strained through 70μm filters. T cells were depleted by staining samples with the following biotinylated antibodies at 1:400 dilution: TCRβ, TCRγδ, CD3ε, CD4, and CD8, then addition of streptavidin microbeads. The cells were passed over LD columns magnetic columns (Miltenyi Cat. #130-042-901). T cell depleted bone marrow cells were resuspended to a final concentration of 10^6^ cells/100μL in ice cold 1X PBS+0.5% FBS. Recipient mice were sublethally irradiated (750 rads or 550 rads) approximately 4 hours before retro-orbital injection with 10^6^ cells. Mice were monitored daily for leukemia for up to 30 weeks post-transplant.

### Cell Sorting for PCR

DN3, DP, and CD4 SP thymocytes were sorted from VcKO, LcKO, and control mice. DNA was isolated using Zymo Research Quick-DNA miniprep kit (Cat. #D3024) following manufacturers protocols. DNA was PCR amplified using the following primers: E2a floxed allele 5’–TCGTCCTCGTCCTCGTCT–3’ E2a deleted allele 5’– CTCACAGAGACCTCCCGACT–3’ and Universal Reverse primer 5’– CGGATCCATCCTCGTCTTTGGTACTG–3’. Standard was made by mixing WT DP DNA and VcKO DP DNA at indicated ratios.

### RNA isolation

CD45.1+ DP thymocytes were isolated from *Vav^Cre^E2a^f/f^*, *Lck^Cre^E2a^f/f^*, *Vav^Cre^E2a^f/f^* + WT BM chimeras, or Ctrl mice via FACS sorting using a FACSAriaIII or FACSAria Fusion cell sorter (BD Biosciences). DPs from *Vav^Cre^E2a^f/f^* and *Lck^Cre^E2a^f/f^* mice were identified as Propidium Iodide^-^CD4^+^CD8^+^ and DPs from *Vav^Cre^E2a^f/f^* + WT BM mice were identified as Propidium Iodide^-^ CD4^+^CD8^+^CD45.1^+^. Cells were sorted into FACS buffer + 1mM EDTA, centrifuged at 400g for 5 minutes, and resuspended in RNeasy lysis buffer. RNA isolation was performed using RNeasy mini kits (QIAGEN) following the manufacturer’s protocol.

### RNA-seq

RNA-seq libraries were constructed using Nugen’s Ovation Ultralow Library systems followed by 100 cycles of sequencing on a NovaSeq 6000 in the Genomics Facility at the University of Chicago. Raw sequence reads were trimmed using Trimmomatic v0.33 (Bolger et al., 2014), and then aligned to the mm10 genome assembly from iGenomes (https://support.illumina.com/sequencing/sequencing_software/igenome.html) using STAR (Dobin et al., 2013). Aligned reads were assigned to genes using the htseq-count tool from HTSeq v 0.6.1 and gene annotations from Ensembl release 78 (Anders et al., 2015). DESeq2 (Love et al., 2014) was used to normalize the gene counts and to calculate differential expression statistics for each gene for each pairwise comparison of sample groups within an nf-Core Differential Abundance workflow (Manning et al., 2023). Metascape analysis was performed on differential gene expression lists (Zhou et al., 2019). Gene set enrichment analysis (GSEA) was performed using gene sets from the Hallmark Pathways of MSigDB (Subramanian et al., 2005). Genes were considered differentially expressed if the Log2FC was > 0.5 had an p-value of <0.05. Data are available through the Gene Expression Omnibus ().

### Statistical analysis

EdgeR or GraphPad Prism software was used to calculate statistics. Unless otherwise noted, a Student’s t-test was used to establish the level of significance between two groups. Groups of 3 or more were assessed using ANOVA with multiple comparisons. Kaplan-Meier curves were analyzed using Log-Rank (Mantel-Cox) test to determine significance. *P<0.05, **P<0.01, ***P<0.001, ****P<0.0001.

## ACKNOWLEDGEMENTS

We thank members of the Kee laboratory, S. Dias, and A. Melnick for helpful discussions and S. Liang for technical assistance. This work was funded by the National Institutes of Health grants R21 AI119894, R01 AI107213, the Janet Rowley Fund from the University of Chicago Comprehensive Cancer Center, and a Team Science award from the Biological Science Division of the University of Chicago to B. L. Kee. G. Parriott was supported by T32 AI007090. We thank the University of Chicago Genomics Facility (RRID: SCR_019196) and Cytometry and Antibody Technology Facility (RRID: SCR_017760), which receive financial support from the Cancer Center Support Grant (P30 CA014599), and the Animal Resource Center.

The authors declare no competition financial interests.

## Author contributions

G. Parriott designed, performed, and interpreted experiments and analyzed RNA-seq data, E. Hegermiller, R.E. Morman and C. Frank maintained mice, performed and analyzed experiments, and had essential intellectual input, C. Saygin and W. Stock provided valuable advice and edited the manuscript, E. T. Bartom performed RNA-seq analysis and edited the manuscript, B. L. Kee conceived of the project, designed, and interpreted experiments and provided funding for the project. G. Parriott and B.L. Kee wrote the manuscript.

## SUPPLEMENTAL FIGURES AND LEGENDS

**Supplemental Fig. 1.**
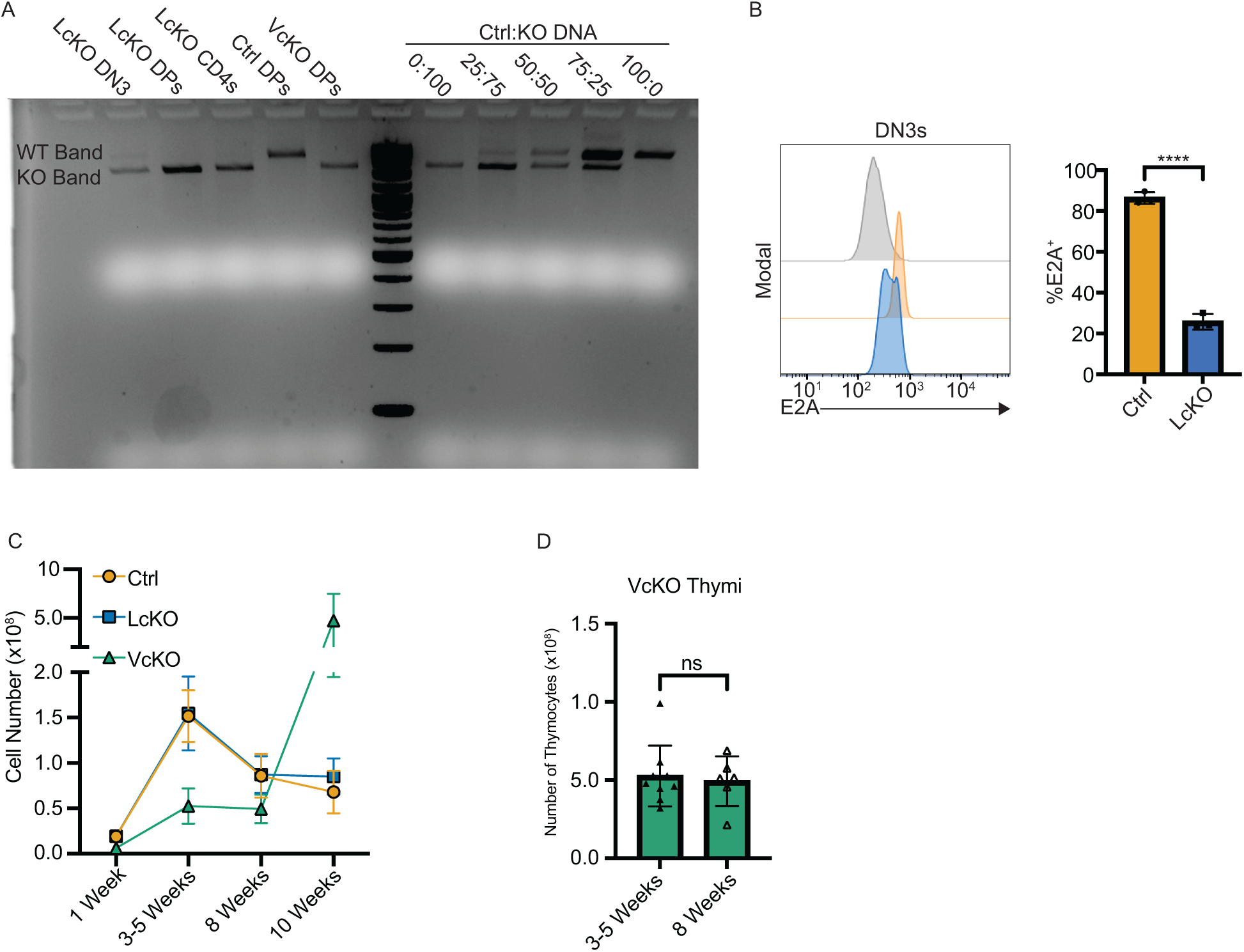
*E2a* is deleted in DN3 thymocytes from LcKO mice. (A) PCR analysis of showing the *E2a*-floxed and *E2a*-deleted alleles in VcKO, LcKO and control thymocytes. (B) Representative FACS histograms and quantification of E2a expression in LcKO and control DN3s. Grey histogram represents LcKO DN3s unstained control. (C) Number of thymocytes from VcKO, LcKO, and control mice over time. (D) Number of thymocytes from 3–5-week-old and 8-week-old VcKO mice.

**Supplemental Fig. 2.**
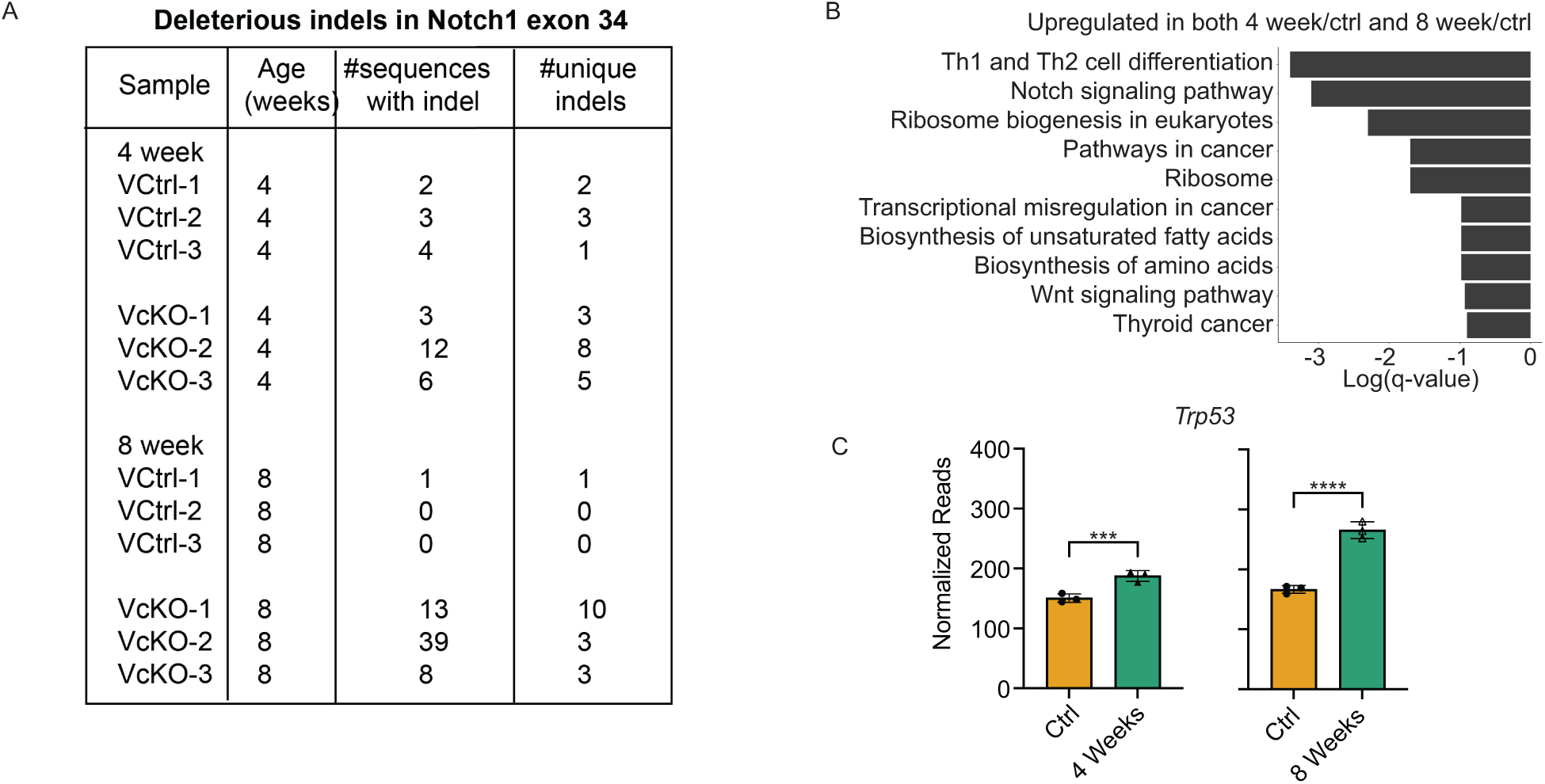
Gene dysregulation in VcKO at 4 and 8 weeks of age. (A) Deleterious indels identified in *Notch1* exon 34 by RNA-seq of DP thymocytes at 4 and 8 weeks of age. (B) Graph of log(q-value) of relevant KEGG pathways analysis of genes differentially expressed in both 4- and 8-week-old VcKO DPs compared to Ctrl DPs. (B) Normalized RNA-sequencing reads showing altered *Trp53* expression in 4-week and 8-week VcKO DPs vs Ctrl. Significance determined using EdgeR (see methods).

**Supplemental Figure 3.**
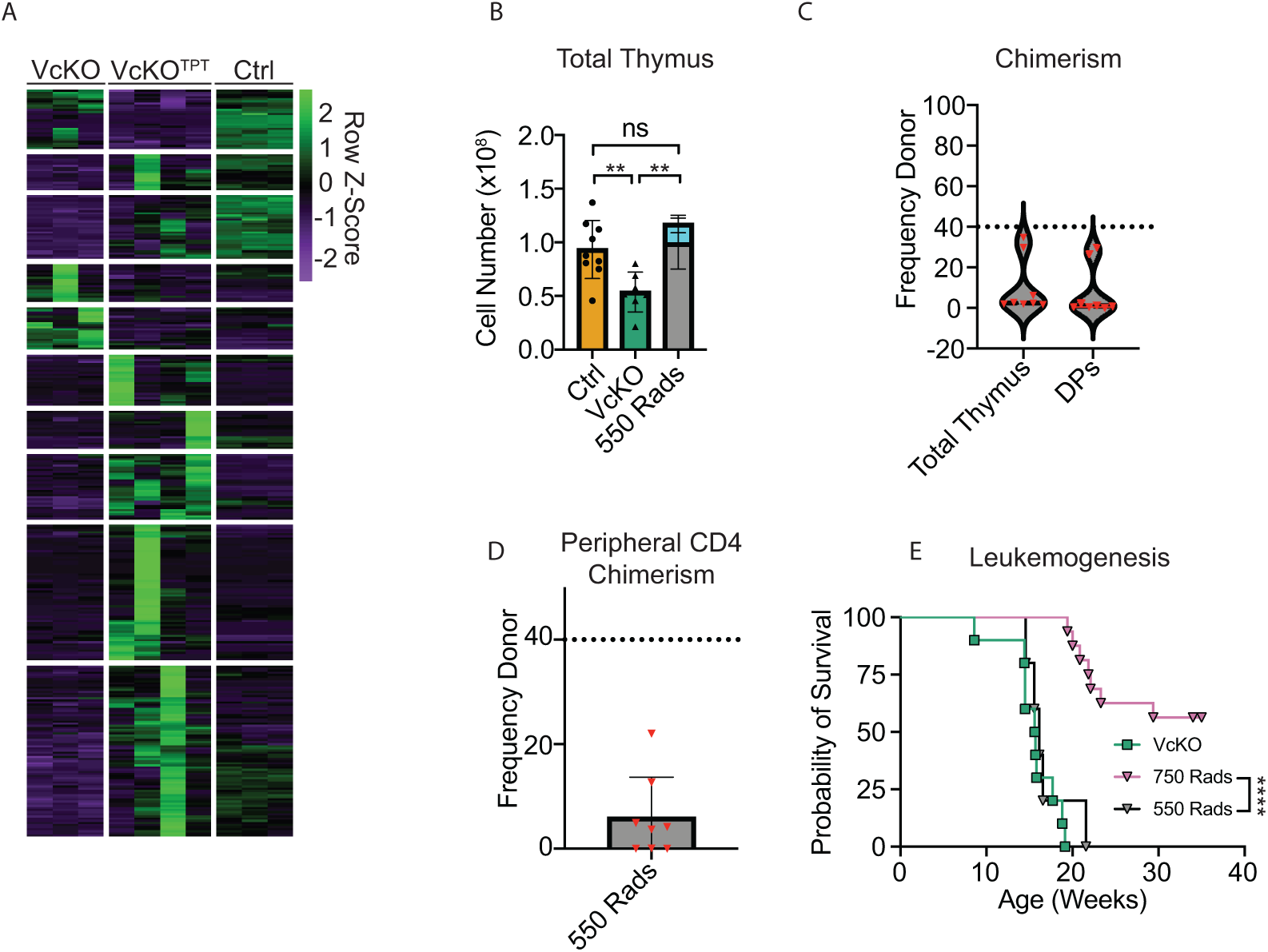
High chimerism is required for competition. (A) Heatmap of DEGs in either VcKO or VcKO^TPT^ DPs compared to Ctrl DPs. (B) Number thymocytes in VcKO, 550Rad VcKO+WT chimeras, and Ctrl mice 4 weeks post-transplant. (C) Graph showing the frequency of donor thymocytes in total thymus and DP population in 550Rad VcKO+WT mice. (D) Graph showing the frequency of donor thymocytes in the peripheral CD4 compartment of 550Rad VcKO+WT chimeras. (E) Kaplan-Meier curve showing survival of VcKO, 750Rad VcKO+WT chimeric, and 550Rad VcKO+WT chimeric mice.

